# Genome Wide Interaction Studies Identify Sex-Specific Risk Alleles for Nonsyndromic Orofacial Clefts

**DOI:** 10.1101/329599

**Authors:** Jenna C. Carlson, Nichole L. Nidey, Azeez Butali, Carmen J. Buxo, Kaare Christensen, Frederic W-D Deleyiannis, Jacqueline T. Hecht, L. Leigh Field, Lina M. Moreno-Uribe, Ieda M. Orioli, Fernando A. Poletta, Carmencita Padilla, Alexandre R. Vieira, Seth M. Weinberg, George L. Wehby, Eleanor Feingold, Jeffrey C. Murray, Mary L. Marazita, Elizabeth J. Leslie

**Author notes:** JCC and NLD should be considered joint first author. Corresponding Author: Elizabeth J. Leslie.

## Abstract

Nonsyndromic cleft lip with or without cleft palate (NSCL/P) is the most common craniofacial birth defect in humans and is notable for its apparent sexual dimorphism where approximately twice as many males are affected as females. The sources of this disparity are largely unknown, but interactions between genetic and sex effects are likely contributors. We examined gene-by-sex (G x S) interactions in a worldwide sample of 2,142 NSCL/P cases and 1,700 controls recruited from 13 countries. First, we performed genome-wide joint tests of the genetic (G) and G x S effects genome-wide using logistic regression assuming an additive genetic model and adjusting for 18 principal components of ancestry. We further interrogated loci with suggestive results from the joint test (p < 1.00 × 10^−5^) by examining the G x S effects from the same model. Out of the 133 loci with suggestive results (p < 1.00 × 10^−5^) for the joint test, we observed one genome-wide significant G x S effect in the 10q21 locus (rs72804706; p = 6.69 × 10^−9^; OR = 2.62 [1.89, 3.62]) and 16 suggestive G x S effects. At the intergenic 10q21 locus, the risk of NSCL/P is estimated to increase with additional copies of the minor allele for females, but the opposite effect for males. Our observation that the impact of genetic variants on NSCL/P risk differs for males and females may further our understanding of the genetic architecture of NSCL/P and the sex differences underlying clefts and other birth defects.

## Introduction

Orofacial clefts (OFCs) are one of the most commonly occurring congenital defects worldwide, affecting 1 in 700 births (Leslie & Marazita, 2013). Incidence rates are vary across populations; to the best of our knowledge, populations with African ancestry have the lowest rates among live births (∼1/2500), European populations have intermediate rates (∼1/1000), and Asian populations have the highest rates (∼1/500) (Mossey, 2007). OFCs are classified as syndromic or nonsyndromic based on the presence or absence, respectively, of additional structural or cognitive abnormalities (Dixon et al., 2011). OFCs are highly heritable (50-80%) based on concordance rates in monozygotic twins (Grosen et al., 2011). This suggests a strong genetic component to the etiology of OFCs.

Approximately 70% of OFCs are considered nonsyndromic and have a multifactorial etiology where genetics, environmental exposures, and their interactions have been shown to modify risk. There have been nine independent genome wide association studies (GWAS) and two meta-analyses that have identified at least 25 genetic loci that contribute to risk for OFCs [(Beaty et al., 2010; Beaty et al., 2011; Beaty et al., 2013; Birnbaum et al., 2009; Grant et al., 2009; Leslie et al., 2017; Leslie, Carlson, et al., 2016; Leslie, Liu, et al., 2016; Ludwig et al., 2012; Sun et al., 2015; Wolf et al., 2015; Yu et al., 2017)]. GWAS has been a useful approach for identifying genetic risk factors for OFCs, however, these studies cumulatively explain only 25-30% of the heritable risk of OFCs in any population (Leslie, Carlson, et al., 2016; Yu et al., 2017).

A likely explanation for the missing heritability of OFCs are non-additive effects, which include interactions with environmental exposures or other genes. Environmental risk factors include maternal tobacco use (M. Shi et al., 2008), low maternal folic acid (Yazdy et al., 2007), and maternal alcohol use. Incorporation of these environmental exposures into genetic studies revealed several gene-environment interactions. These include a 3-to 6-fold increased risk for OFCs for fetuses with null alleles of glutathione s-transferases (*GSTT1* and *GSTM1*) and whose mothers smoked during pregnancy (Lammer et al., 2005; Min Shi et al.). In genome-wide studies, several gene-environment (G x E) interactions have been identified (Wu et al., 2010; Wu et al., 2014). Until recently, G x E interactions were the only genome-wide significant association signals for cleft palate (Beaty et al., 2011; Leslie, Liu, et al., 2016).

An understudied factor has been the contribution of sex to OFC risk. Males are twice as likely to have CL or CLP than females (Marazita, 2012) and females are twice as likely to have CP than males, which implies that cleft type and incidence are sexually dimorphic. Explanations for these incidence rates have been elusive, but several genetic models could explain the reduced risk for CL/P in females. First, there may be specific susceptibility factors on the X or Y chromosomes that affect male-CL/P risk, but not female-CL/P risk, due to the compensatory second copy of the X chromosome or lack of a Y chromosome in females. There are several notable candidate gene association signals on the X-chromosome, including *TBX22*, associated with CP (Marcano et al., 2004; Suphapeetiporn et al., 2007), and *EFNB1* and *DMD* (Patel et al., 2013). Several other loci have been identified using a variety of methods and genetic models (Jugessur et al., 2012; Patel et al., 2013; Skare et al., 2017). X-chromosome inactivation has been investigated in female discordant CL/P twin and sibling pairs, suggesting a role for this mechanism in CL/P (Kimani et al., 2007). Second, there may be different liability thresholds in which the same risk alleles affect males and females equally, but females have a higher threshold that requires a higher load of risk alleles or highly penetrant mutations. Finally, risk factors could be located on the autosomes with differential effects in males and females, perhaps due to hormonally-mediated dimorphism. The third model is the focus of the present study as such gene-by-sex (G x S) interactions have not yet been directly tested.

It is well established that a standard case-control analysis has low power to detect interactions because of small numbers of cases or controls when stratified by the exposure. Alternative approaches have been developed for testing interactions in the context of genome-wide association analyses. We elected to perform a two-step procedure to gain power by enriching for possible G x S interactions by first jointly testing for genetic main effect and G x S interaction (Dai et al., 2012; Kraft et al., 2007). We therefore sought to determine how sex contributes to OFC sex-specific risk with a genome-wide analysis of G x S interactions, and potentially to identify new OFC risk loci, using our large, multi-ethnic OFC population.

## Methods

### Study Population

The subjects for this study were drawn from our large, multi-ethnic OFC cohort we described previously (Leslie, Carlson, et al., 2016). We selected 5,984 subjects from 18 recruitment sites, located in Africa, Asia, Latin America and North America (Table 1). The recruitment sites were a part of genetic studies being conducted by the University of Iowa and University of Pittsburgh Center for Craniofacial and Dental Genetics. All sites had local Institutional Review Board approval and subjects were recruited using informed consent procedures. All subjects consented to the use of their DNA in genomic studies. Cases were defined as unrelated individuals with nonsyndromic cleft lip with or without cleft palate (CL/P). Controls were defined as unrelated subjects with no personal or family history of any structural or developmental anomaly. Sex was confirmed genetically for all subjects as part of quality control procedures.

**Table 1:** Study Population

| **Cleft Type** | **Males** | **Females** | **Total** |
| --- | --- | --- | --- |
| CL/P | 1293 | 849 | 2142 |
| CL | 251 | 199 | 450 |
| CLP | 1042 | 650 | 1692 |
| Controls | 713 | 987 | 1700 |

### Genotyping and quality control

The methods for genotyping, quality control, imputation, and deriving principal components (PCs) of ancestry have been previously described (Leslie, Carlson, et al., 2016). Briefly, samples were genotyped for ∼580K SNPs at the Center for Inherited Research at John Hopkins University using the Illumina HumanCore+Exome chip. Approximately 97% of the genotyped SNP’s passed quality control leaving a total of 539,473 genotyped SNPs. Low minor allele frequency (MAF <5%) SNPs were excluded from analysis. IMPUTE2 software was utilized for imputation of 34,985,077 unobserved polymorphisms using the multiethnic 1000 Genomes Project Phase 3 reference panel.

Imputed genotype probabilities were converted to most-likely genotypes using GTOOL; only most-likely genotypes with probabilities > 0.9 were retained for statistical analysis. Imputed SNPs with INFO scores < 0.5 or those deviating from Hardy-Weinberg equilibrium (p < 1.00 × 10^−4^ in European controls) were also excluded from the analysis. Population structure was investigated by using a principal components analysis using the R package SNPRelate following the approach introduced by Patterson et al. (2006). The University of Washington Genetics Coordinating Center released a full online report on the data cleaning, quality assurance, ancestry analyses and imputation for this study (http://www.ccdg.pitt.edu/docs/Marazita—ofc—QC—report—feb2015.pdf, last accessed April 25, 2016).

### Statistical Analysis

We examined potential gene-by-sex interactions following a case-control approach first introduced by Kraft et. al (Kraft et al., 2007). In the first step, logistic regression examining main effects of genotype (using the additive genetic model) and sex, as well as the interaction between them, was performed using PLINK (v1.9) (Purcell et al., 2007). Eighteen principal components of ancestry were identified and used as covariates in the analysis. A two degree of freedom likelihood ratio test was used to assess the joint association of the genetic and gene-by-sex components. SNPs demonstrating at least moderate evidence of association (i.e. p-value < 1.00 × 10^−5^) in the two degree of freedom test were then individually examined in the second step. Because there are very strong main effects of genotype for CL/P, we then interrogated the G x S term (from the same model) to identify significant interactions, using a genome-wide significance threshold of 5.00 × 10^−8^ and a suggestive threshold of 1.00 × 10^−5^. Overall, this approach examines potential genetic risk factors for OFC which differ between males and females.

## Results

We previously performed conventional tests of association to identify marginal gene (G) effects increasing risk for NSCL/P (Leslie, Carlson, et al., 2016). Here we carried out a genome-wide screen for G x S interaction where we first examined the 2df test for gene (G) and gene-sex (G x S) interaction. Our analysis identified four loci that achieved genome-wide significance in the 2df test. Three of these loci (*IRF6* on 1q32, 8q24, and *NTN1* on 17p13) showed very strong G effects, consistent with our previous analysis of this dataset (Leslie, Carlson, et al., 2016). The fourth locus, on 10q21, represented a new signal (lead SNP rs72804706, p_2df_ = 3.67 × 10^−8^), driven by a strong G x S interaction.

We then examined the test for G x S interaction alone for all SNPs demonstrating at least suggestive evidence of association in the 2df test. Because the goal of this study was to identify interactions, we focused on signals showing evidence of G x S interaction (Figure 1). 128 loci containing 1,649 SNPs surpassed a suggestive p-value threshold in the 2df test. Of these, 252 SNPs from 17 loci showed evidence of a suggestive statistical G x S interaction (p_GxS_ < 1.00 × 10^−5^), with one locus demonstrating a genome-wide significant interaction effect (10q21, lead SNP rs72804706, p_GxS_ = 6.69 × 10^−9^; Table 2). Interestingly, we did not find evidence of sex-specific effects for any previously identified NSCL/P risk loci.

**Table 2.**
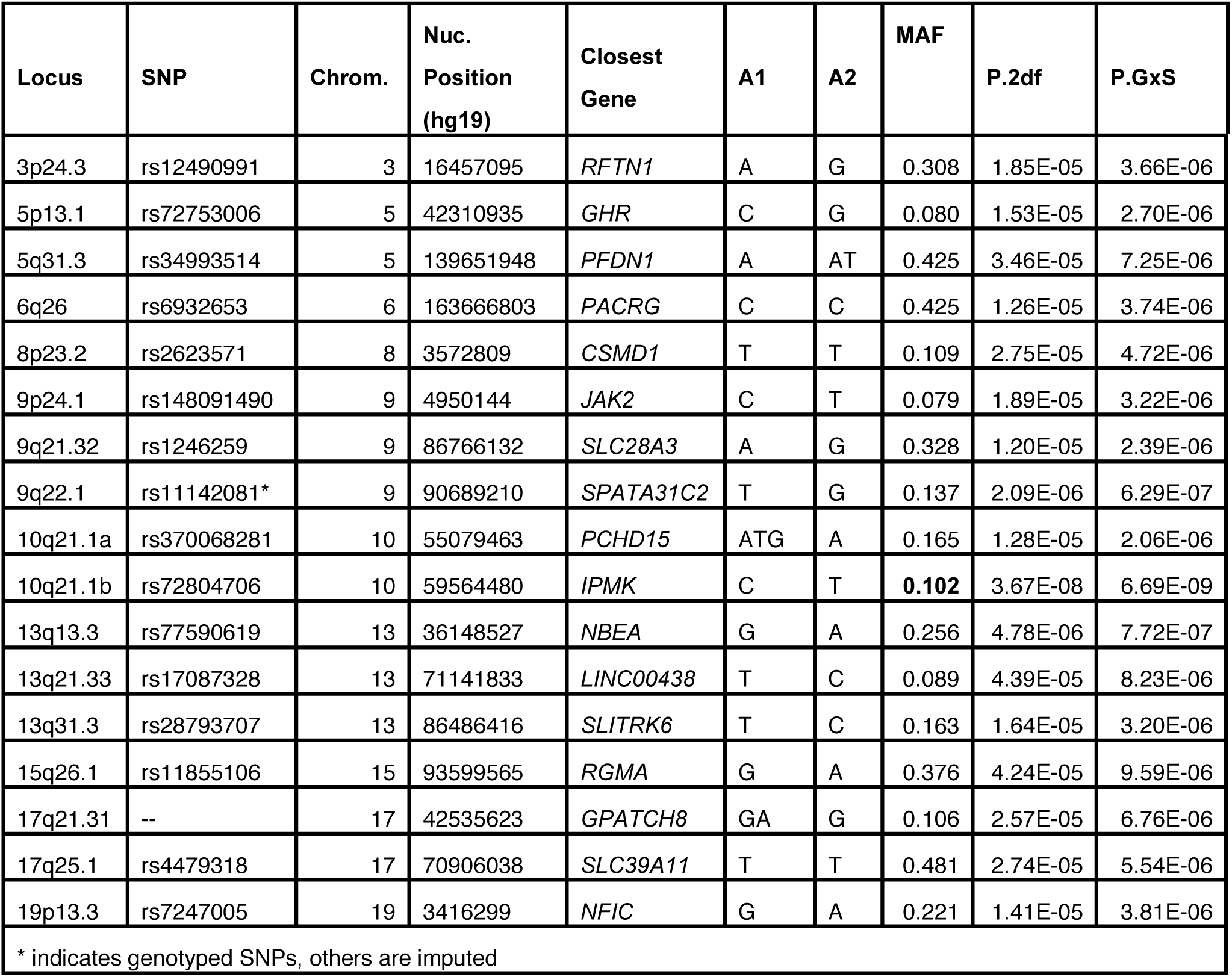
17 loci with evidence for G x S interactions

**Figure 1.**
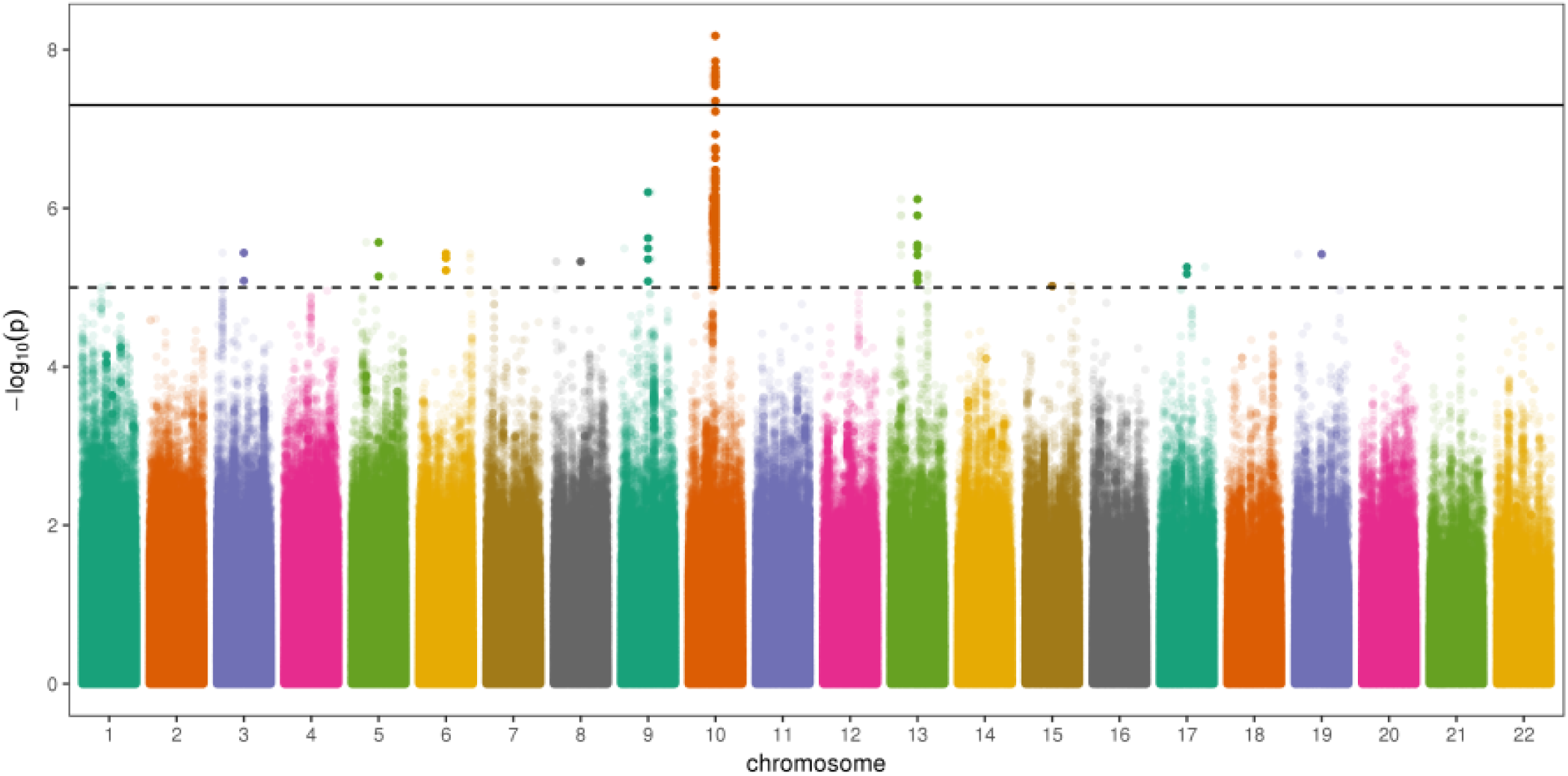
Manhattan plot displaying the p-values from the gene-by-sex effect GWAS. Bolded points indicate a variant for which p-value for the two degree of freedom joint test of gene and gene-by-sex effect was less than 1.00 × 10^−5^.

The 10q21 locus included 220 SNPs with evidence of G x S interaction (Figure 2B). For lead SNP rs72804706, we estimated the odds of CL/P to be 28% lower in males for each additional copy of the T allele. By contrast, the odds of CL/P in females were estimated to be 81% higher with each additional copy. However, it is worth noting that males had much higher estimated risk of CL/P than females for individuals with the homozygous reference genotype; the interaction effect is realized in the increasing risk for females with copies of the risk allele, T (Figure 2E). Post-hoc power was estimated to be 75% for the effects observed with rs72804706 (Kooperberg, 2015).

**Figure 2.**
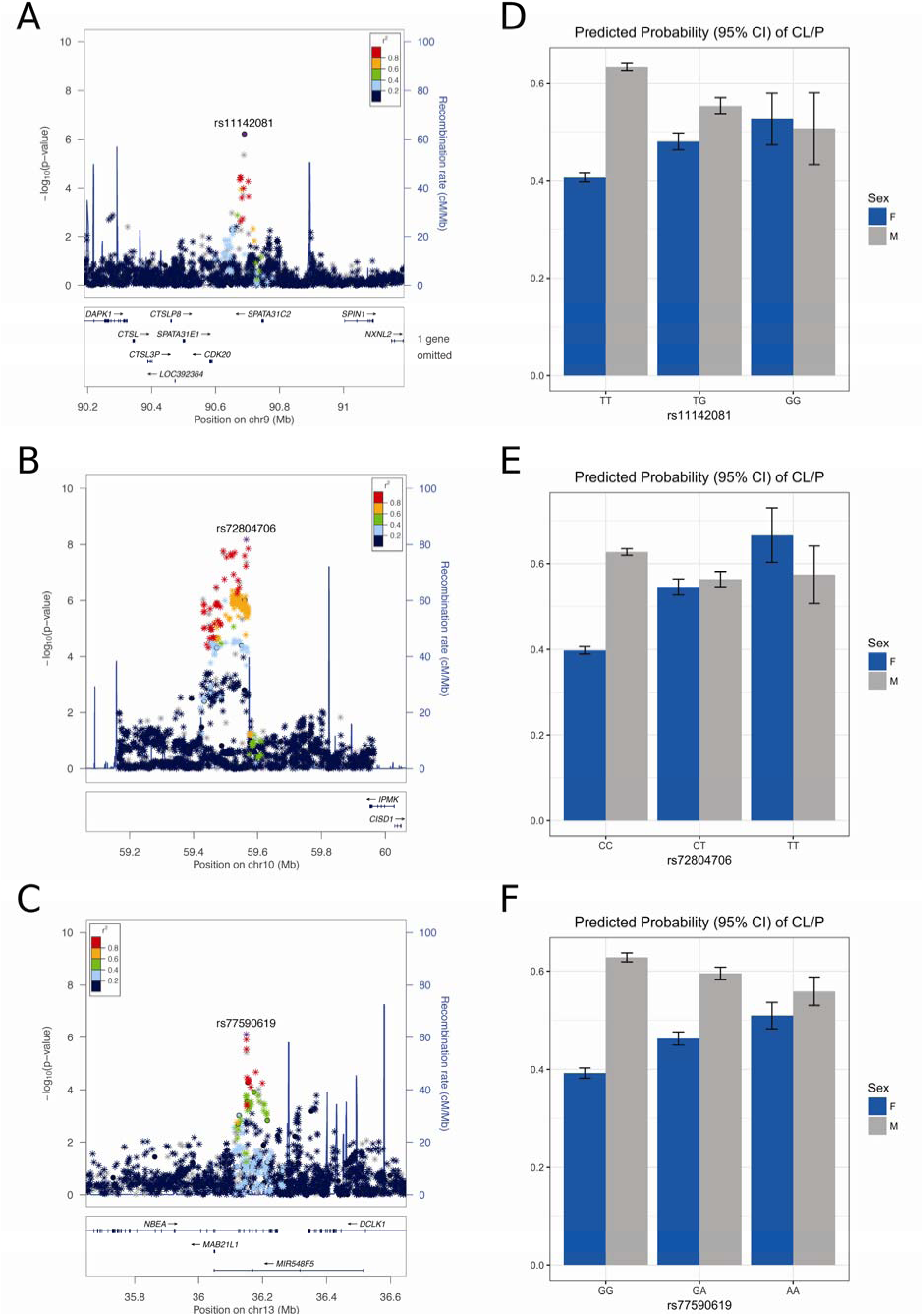
Results for the 9q22.1 (A,D), 10q21.1 (B,E), and 13q13.3 (C,F) loci. Regional association plot show results for the gene-by-sex effect model. Points are color-coded based on linkage disequilbrium (r^2^) in Europeans. Plots were generated with LocusZoom (Pruim et al., 2010). Bar charts of the predicted probability of CL/P are shown, stratified by sex and adjusting for principal components of ancestry. Plots were generated with the ggplot2 package in R (R Core Team, 2017; Wickham, 2009).

Sixteen loci had suggestive evidence of G x S interactions, including loci on 9q22.1 and 13q13.3 with G x S interactions approaching genome-wide significance (p < 1.00 x 10^−7^; Figure 2A,C). At 9q22.1, for lead SNP rs11142081, we estimated the odds of CL/P to be 28% higher in females and 31% lower in males for each additional copy of the G allele (Figure 2D). At 13q13.3, for lead SNP rs77590619, the odds of CL/P were estimated to be 28% higher in females and 21% lower in males for each copy of the A allele (Figure 2F). The findings for these loci are similar to that for 10q21, in that males had a much higher risk of CL/P for individuals with the reference homozygous genotype.

Because the incidence of CL/P among males and females is opposite the incidence of CP (CP being more common among females), we next wanted to determine if the direction of the effect for our top SNPs was the same among CP cases. Because most genetic studies have focused on CL/P, the samples sizes are still quite small for CP (Leslie, Liu, et al., 2016). As interaction tests require much larger samples sizes to maintain statistical power than tests for main effects alone, we performed a qualitative analysis of the unadjusted minor allele frequencies of these SNPs in CP cases and controls (Table 3). The frequencies of the lead SNP at 10q21 did not suggest an interaction for CP, but the frequencies of 13q13.3 and 9q22.1 SNPs were consistent with the trends we observed in the CL/P analysis (higher frequencies among female cases and male controls).

**Table 3.**
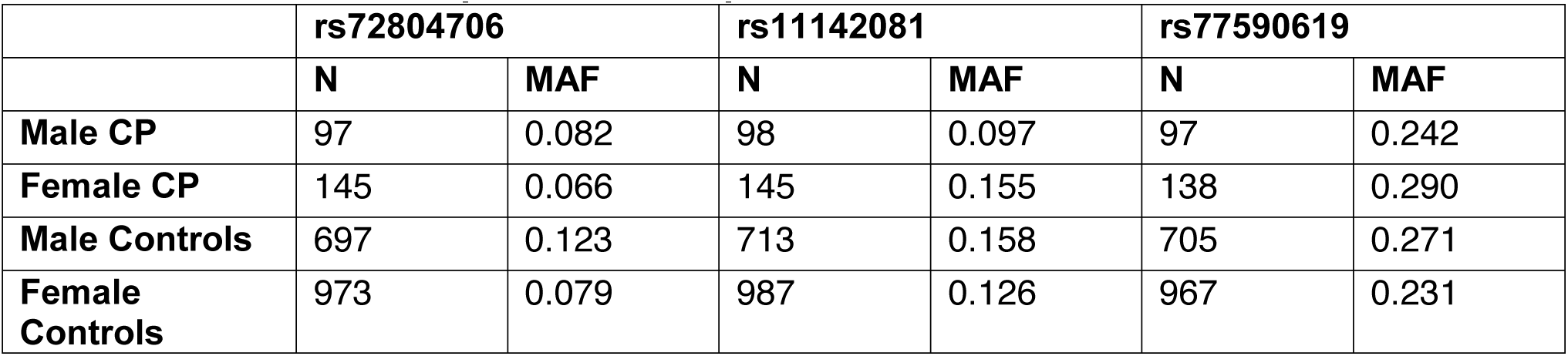
Minor allele frequencies for top CL/P loci in CP cases

## Discussion

Our results support interaction between SNPs in autosomal genes and sex leading to higher risk of CL/P. This finding is not surprising as incidence rates of CL/P differ by sex, however, this is the first study to use a genome-wide approach to identify the genetic contribution to this phenomenon. Outside of the OFC literature, there is substantial evidence that genetic variation has sex-dependent effects; a 2014 study catalogued 33 sex-dependent loci in 26 medically-relevant traits (Gilks et al., 2014). The well-documented advantages of the unbiased GWAS are useful in studies such as ours, where the mechanisms of the CL/P male bias are not understood. Consequently, this study should be viewed as hypothesis generating because gene-by-sex interactions identify modifiers whose biological impact could occur far upstream of craniofacial development. Model systems provide evidence for some of these mechanisms, such as sex-specific trans-eQTLs (Derome et al., 2008) and sex-specific epistasis (Bhasin et al., 2008), but the biochemical causes of these effects likely involve sex hormones, sex-biased methylation (McCarthy et al., 2014), and/or interactions with sex chromosomes. Given the diversity of possible mechanisms, it is impossible to adequately explore or discuss each locus, so we will only briefly discuss what is known about the genes located closest to the top three association signals.

The primary finding of this study was a novel locus at 10q21.1 with a genome-wide significant G x S interaction for multiple SNPs. The lead SNP, rs72804706, is located in a large intergenic region 400kb upstream of inositol polyphosphate multikinase (*IPMK*). Within 500kb either side of the lead SNP are formally significant and suggestive association signals from genome-wide association studies covering a wide range of traits and disease including diabetic retinopathy, aggressiveness in attention deficit hyperactivity disorder, alcohol dependence, cardiovascular traits, and inflammatory bowel disease (MacArthur et al., 2017). Further, a truncation mutation in IPMK was identified by linkage and exome sequencing in a large family with small intestinal carcinoids (Sei et al., 2015). Although it is not known if any of the associated nearby SNPs regulate *IPMK*, the variety of disease process is consistent with the diverse roles of IPMK. IPMK is considered to be a pleiotropic protein with both kinase and non-catalytic functions in multiple cellular processes, including transcriptional coactivation, corepression, and cell signaling (Kim et al., 2017; Xu et al., 2013). *IPMK*-null mice die early in embryonic development and have multiple morphological defects, including neural tube misfolding (Frederick et al., 2005). Tissue-specific knockouts will be needed to dissect the specific roles of IPMK in OFCs and other disease.

Among our other findings are loci at 13q13.3, and 9q22.1. The 13q13.3 lead SNP is located within an intron of neurobeachin (*NBEA*), a synaptic scaffolding protein associated with autism (Castermans et al., 2003) and migraine in bipolar disorder (Jacobsen et al., 2015). Little is known about spermatogenesis-associated protein 31C2 (*SPATA31C2*), the proximal gene at the 9q22.1 locus. The implicated variants from our analyses in these three regions are associated with higher risk of CL/P for females carrying the minor risk allele; this trend is not present in males, who have higher incidence of CL/P than females.

We did not perform a formal interaction analysis for CP, but found a similar trend in unadjusted minor allele frequencies for our associated SNPs for the 9q22.1 and 13q13.3 loci. This would suggest that these loci we identified do not completely explain the bias in CL/P and CP incidence in males and females because if they did, we would expect no effect or the opposite effect in CP. Thus, the alleles at these loci may increase risk for any kind of OFC in sex-specific manner. Although the female palate is known to close approximately one week later than the male palate, increasing risk for CP and possibly explaining the female preponderance of CP (Burdi & Silvey, 1969), genetic variants may contribute to this delay so formal interaction studies should be performed in this OFC subtype. Altogether our current findings demonstrate the capability of genome-wide interaction studies to identify sex-specific effects, but more research is warranted to investigate the underlying sex discrepancy in the incidence of both CL/P and CP.

## Acknowledgments

We gratefully acknowledge the participation of the families, field staff, and collaborators around the world who made this project possible. This work was supported by grants from the National Institutes of Health (NIH): R00-DE025060 [EJL], X01-HG007485 [MLM, EF], R01-DE016148 [MLM, SMW], U01-DE024425 [MLM], R37-DE008559 [JCM, MLM], R01-DE009886 [MLM], R21-DE016930 [MLM], R01-DE014667 [LM], R01-DE012472 [MLM], R01-DE011931 [JTH], R01-DE011948 [KC], U01-DD000295 [GLW], K99-DE024571 [CJB]; and PICT-2016-3869 [FAP].], S21-MD001830 [CJB], U54-MD007587 [CJB]). Additional support was provided by the Philippine Band of Mercy, the Hope Foundation Inc., Operation Smile, and the Noordhoff Craniofacial Foundation Philippines Inc. Genotyping and data cleaning were provided via an NIH contract to the Johns Hopkins Center for Inherited Disease Research: HHSN268201200008I.

## Grant Numbers

R00-DE025060, X01-HG007485, R01-DE016148, U01-DE024425, R37-DE008559, R01-DE009886, R21-DE016930, R01-DE014667, R01-DE012472, R01-DE011931, R01-DE011948, U01-DD000295, K99-DE024571; PICT-2016-3869, S21-MD001830, U54-MD007587

